# *Rlim* coordinates diurnal regulation of food intake and thermogenesis

**DOI:** 10.1101/2025.07.23.666379

**Authors:** Feng Wang, Poonam Mehta, Max Zinter, Peter M’Angale, Timmy Lê, Huawei Li, Jason K. Kim, David A. Guertin, Gilles E. Martin, Travis Thomson, Ingolf Bach

## Abstract

Energy homeostasis in mice is maintained through coordinated activity among hypothalamic nuclei that regulate food intake and thermogenesis. These processes must adapt to the sleep–wake cycle, yet the underlying pathways, cell types, and molecular mechanisms governing their diurnal regulation remain poorly understood. We show that mice lacking the E3 ubiquitin ligase *Rlim* are lean and resistant to diet-induced obesity, owing to reduced food intake and enhanced brown adipose tissue (BAT) thermogenesis. We identify GABAergic neurons in the suprachiasmatic nucleus (SCN)—components of the central circadian clock—as mediators of these effects. Specifically, *Rlim* in RIP-Cre^+^ neurons governs daily thermogenic rhythms, while *Rlim* in vasoactive intestinal peptide (VIP)-expressing neurons modulates diurnal feeding behavior. Thus, *Rlim* is a key regulator of diurnal rhythms controlling energy balance.

## Introduction

The hypothalamus of the brain is composed of several interconnected nuclei that coordinate endocrine functions and maintain body homeostasis. Among these, the arcuate (Arc) and paraventricular (PVN) nuclei are well-established regulators of energy balance, integrating leptin signaling and modulating food intake and brown adipose tissue (BAT) thermogenesis (*1–3*) via GABAergic pathways (*1, 4–7*). Located in the Suprachiasmatic Nucleus (SCN), the circadian clock also influences energy homeostasis (*8–10*), but how these functions are orchestrated remains poorly understood.

The X-linked gene *Rlim* (also known as *Rnf12*) encodes an E3 ubiquitin ligase (*11*) that is broadly expressed, including in the central nervous system (*12*). RLIM protein shuttles between the nucleus and cytoplasm in a phosphorylation-dependent manner predominantly localizing to the nucleus (*13, 14*), where it modulates gene transcription via proteasome-dependent regulation of transcriptional complexes (*11, 15–17*). In female mice *Rlim* is essential for imprinted X chromosome inactivation (iXCI) in placental trophoblasts (*18–20*), and its loss causes early embryonic lethality. Even though male mice systemically lacking *Rlim* are viable and fertile (*18, 20, 21*), *Rlim* contributes to the cytoplasmic recycling during spermiogenesis (*22*). These diverse sex-specific functions are predicted to impact the energy balance in animals.

In this study, we show that mice lacking *Rlim* are lean due to lower food intake and increased thermogenesis in BAT. We find that *Rlim* coordinates diurnal rhythms of BAT thermogenesis and feeding behavior. This is carried out by *Rlim* activity in specific GABAergic neuronal subpopulations characterized by expression of RIP-Cre or vasoactive intestinal peptide (VIP), located in the SCN and participating in the central circadian clock.

## Results

### Mice systemically lacking Rlim are lean and resistant to diet-induced obesity

We found that males systemically lacking *Rlim* exhibited significantly reduced body mass compared to their male littermates carrying a functional *Rlim* allele. Under normal chow diet (ND) at room temperature (RT; ∼21°C), both germline *Rlim* KO/Y and conditional *Rlim* fl/Y; Sox2-Cre⁺/⁻ males gained less weight than controls, despite similar body lengths (Figs. S1A, B). Proton magnetic resonance spectroscopy (1H-MRS) revealed that this weight difference was due to reduced fat mass (Fig. S1C). A similar, though milder, phenotype was observed in females lacking *Rlim* (flm/KOp; Sox2-Cre⁺/⁻), where the maternally inherited floxed allele was targeted after iXCI (*21*) (Fig. S1D). These effects were markedly amplified when mice were switched to a high-fat diet (HFD; fat Calories 60%) at 12 weeks of age (Figs. 1A-C). Indeed, while organ weights including heart, kidney and spleen remained unchanged, *Rlim*-deficient mice had significantly reduced adipose tissue mass—including epididymal (eWAT), inguinal (iWAT), and brown adipose tissue (BAT) (Figs. 1C-E), as well as liver (Figs. 1C; F), exhibiting decreased lipid loads (Figs. 1E, G-I). These findings indicate that *Rlim* deficiency confers protection against diet-induced obesity and hepatic steatosis.

**Fig. 1:**
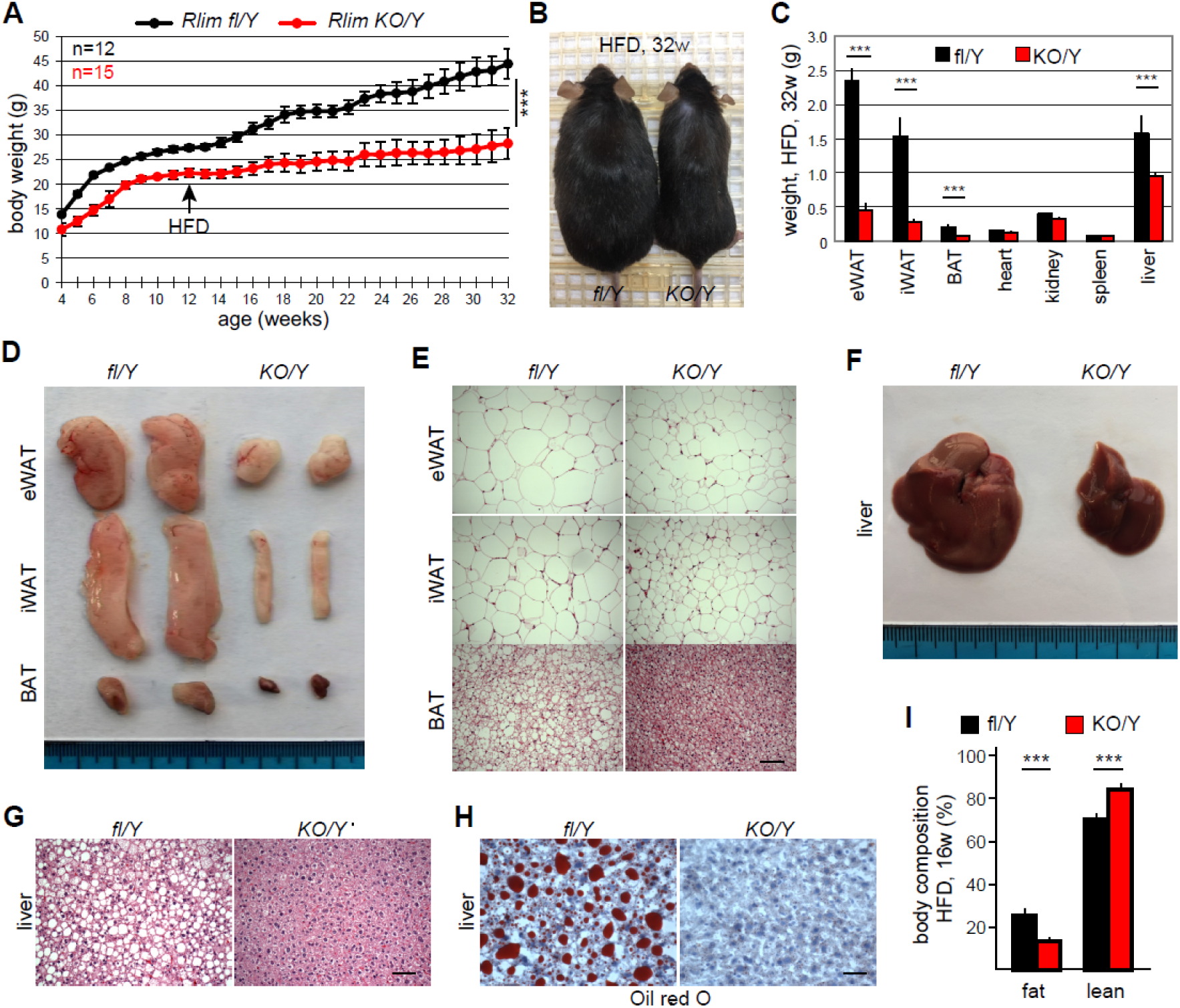
Mice lacking *Rlim* are protected from diet-induced obesity and liver steatosis. Germline *Rlim* KO (KO/Y, red) males and control littermates (fl/Y, black) were fed 12 weeks with ND and then switched over to HFD. *** = P<0.001; Student’s t-test. Error bars indicate SEM. **A)** Animal body weight profiles. Numbers of animals (n) are indicated. **B)** Appearance of animals at 32 weeks. **C)** Comparative organ weights. Note significantly decreased weights of various adipose tissues as well as liver in KO/Y animals. **D)** Representative adipose tissues isolated from animals at 32w. **E)** H&E staining of adipose tissue sections reveals diminished lipid load in droplets of KO/Y animals. Scale bar = 60 μm. **F)** Representative livers of animals at 32w. **G)** Diminished lipid load in liver sections of KO/Y animals as revealed by H&E staining, Scale bar = 60 μm, or **H)** Oil Red O staining, Scale bar = 40 μm. **I)** Decreased relative fat mass and increased relative lean mass in KO animals as measured via 1H-MRS. eWAT=epididymal white adipose tissues, iWAT=inaugural white adipose tissues, BAT= brown adipose tissues.

### Decreased food intake and increased BAT thermogenesis in Rlim-deficient mice

To investigate the basis of the lean phenotype, we assessed food intake and thermogenic activity. *Rlim*-deficient mice consumed significantly less food under both ND and HFD conditions (Fig. 2A), with no evidence of increased fecal energy loss (Fig. 2B). Infrared imaging revealed elevated surface temperatures in scapular BAT regions of *Rlim* KO/Y mice on HFD (Fig. 2C), and in metabolic cages, these mice showed increased energy expenditure without changes in physical activity (Figs. 2D, E). Consistent with enhanced thermogenesis, expression of uncoupling protein 1 (UCP1)—a key thermogenic effector (*23, 24*)—was elevated at both mRNA and protein levels (Figs. 2F, G).

**Fig. 2:**
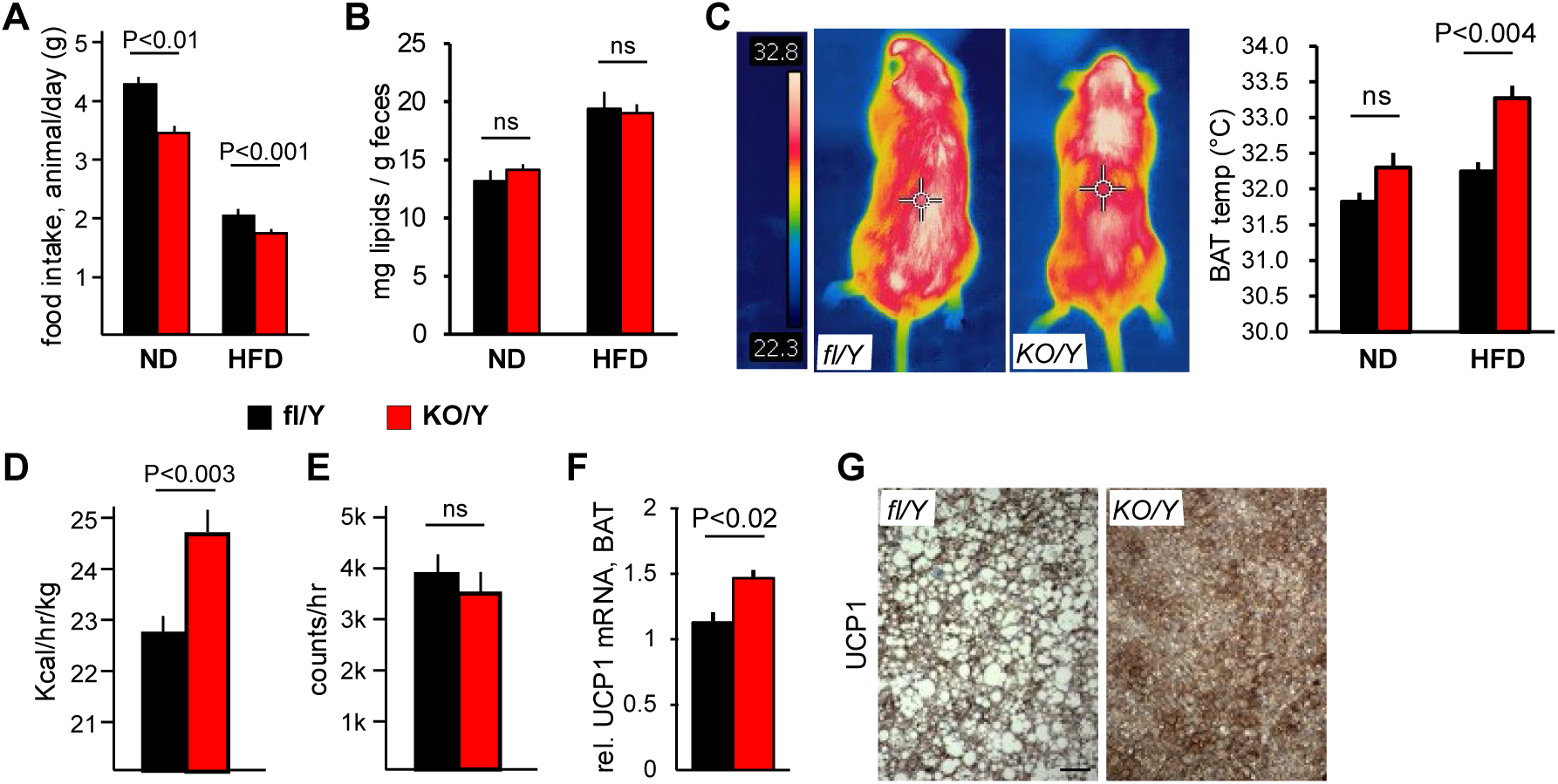
Decreased food intake and increased BAT thermogenesis in *Rlim* KO mice. **Black** = fl/Y; **Red** = KO/Y. P-values: Student’s t-test. Error bars indicate SEM. **A)** Mice lacking *Rlim* eat significantly less than littermate controls. ND=Normal Diet (11 weeks), HFD=High Fat Diet (17 weeks). **B)** Lipid extraction from mouse feces reveals similar lipid content. **C)** Measurements using an infrared camera shows increased thermogenesis in scapular BAT of males systemically lacking *Rlim*. Left: Representative examples of mice fed HFD; right: summary of mice fed ND or HFD. **D, E)** Increased energy expenditure (D) but similar physical activity (E) in mice lacking *Rlim* as measured in metabolic cages (HFD). **F, G)** Increased UCP1 mRNA levels as determined via RT-qPCR (F), and UCP1 protein (IHC) in BAT (G) of KO/Y animals (HFD). Scale bar = 40 μm.

To test whether these effects were developmental in origin or based on a persistent *Rlim* activity, we induced a systemic *Rlim* deletion in adult *Rlim* fl/Y; UBC-Cre/ERT2 mice using tamoxifen. These mice exhibited a partial weight phenotype (Fig. S2), consistent with persistent functions of *Rlim* during the balancing of energy homeostasis. However, due to limited penetrance of tamoxifen-induced expression of Cre in this model, developmental contributions cannot be fully excluded.

### Rlim functions in GABAergic neurons of the hypothalamus

Because of the stereotypical weight gain profile exhibited by mice, a robust weight phenotype (Figs. 1, S1), and efficient *Rlim* deletion across Cre drivers (*18–22, 25*), we used tissue-specific knockouts to identify the relevant tissues and cell types of its action. As *Rlim* is X-linked and males harbor a single maternally transmitted X chromosome, we took advantage of X-linked genetics to target the *Rlim* cKO, mating fl/fl females with males WT for *Rlim* and heterozygous for various Cre drivers (as ordered from The Jackson Laboratories, genotype WT/Y; Cre +/-; Table S1), directly analyzing next generation males. Thus, males that receive the Cre transgene are cKO/Y (**red**), and those that do not serve as fl/Y (**black**) littermate controls. For this mating strategy, Cre-only control experiments were carried out for all drivers that induced a weight phenotype (insets).

Because of the robust phenotype in adipose tissues (Fig. 1), whose functions are partially regulated by the brain (*5*), we targeted the *Rlim* cKO to all adipose tissues via Adiponectin-Cre and to neural tissues via Nestin-Cre (Figs. 3A, S3A). The cKO via Nestin-Cre fully recapitulated the systemic KO phenotype, suggesting that *Rlim* acts in neural tissues. Further dissection using various cell type–specific Cre drivers showed that deletion of *Rlim* in astrocytes (GFAP-Cre), excitatory neurons (CamK2-Cre), cholinergic (Chat-Cre), or dopaminergic neurons (Dat1-Cre) had no effect (Figs. S3B-E), while the deletion in GABAergic neurons (Vgat-Cre) reproduced the lean phenotype (Figs. 3B) and the effects on food intake and BAT thermogenesis (Figs. 3C, D). Because GABA may play roles in other cell types affecting energy homeostasis such as pancreatic β cells (*26*), enteroendocrine cells (EE) of the gut and neurons of the enteric nervous system (ENS) (*27*), we used Cre drivers independently targeting these cell types/tissues including Ins1-Cre (β cells), Pdx1-Cre (pancreatic cell types including β cells), Vil1-Cre (gut epithelial cells including EE cells), and Sox10-Cre (neural crest cells/ENS) (Figs. 3E-G; S3F, see also Table S1). As no weight effect was observed, these results indicate that *Rlim* acts in central Vgat^+^ neurons to regulate energy balance, reminiscent to the situation reported for the leptin pathway (*5*).

**Fig. 3:**
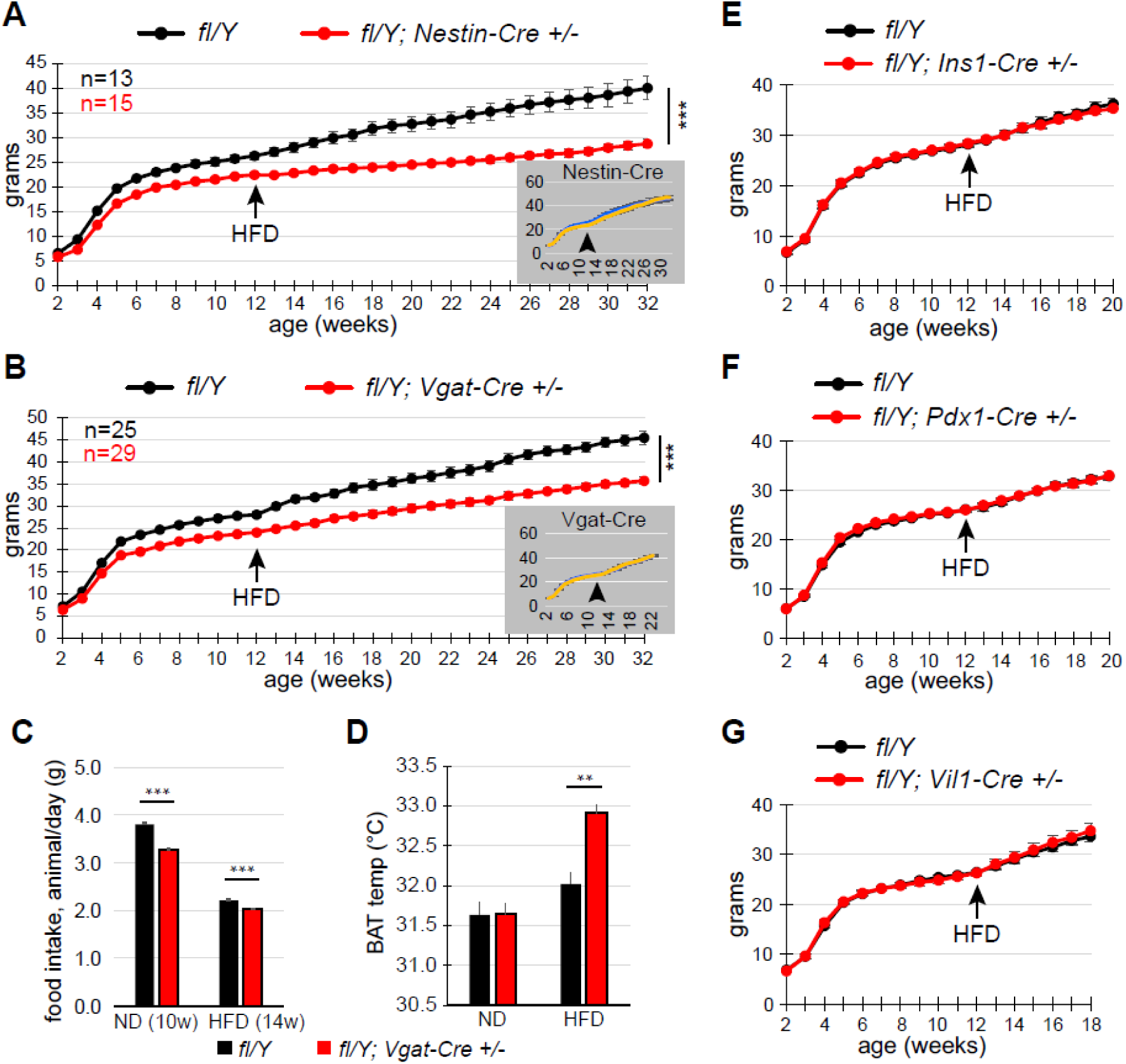
*Rlim* in GABAergic neurons regulates food intake and BAT thermogenesis. Weight profiles: **Black =** fl/Y; **Red** = cKO/Y; Insets: Cre-only controls **blue** = Cre-; **yellow** = Cre^+^). P-values: Student’s t-test; *** = P<0.001; ** = P<0.01; Error bars indicate SEM. **A, B)** Inducing the *Rlim* cKO via Nestin-Cre (A) or Vgat-Cre (B) largely recapitulates weight effects observed in germline KO/Y mice. **C, D)** Food intake (C) and BAT temperature (D) in Vgat-Cre animals. **E)** Targeting of the *Rlim* cKO in pancreatic β-cells via Ins1-Cre, **F)** Pancreatic cell types including β-cells via Pdx1-Cre, **G)** Gut epithelial cell including enteroendocrine cells via Vil1-Cre,

We next focused our studies on neurons of the hypothalamus, where GABAergic neurons mediate much of leptin’s effects on the regulation of food intake and BAT thermogenesis (*5*). Inducing the *Rlim* cKO in hypothalamic VMN or PVN nuclei via SF1-Cre and Sim1-Cre (*28*), respectively, (Figs. S4A, B), or in known neuron subtypes that control energy homeostasis via LepR-Cre, POMC-Cre or AgRP-Cre (Figs. 4A-C) did not affect weight profiles. For testing potential effects in RIP-Cre^+^ neurons, we directly compared *Rlim* WT/Y; RIP-Cre+/-with fl/Y; RIP-Cre +/-animals, thereby controlling for reported effects of this Cre transgene in pancreatic β cells (*29*). Indeed, a RIP-Cre mediated *Rlim* cKO produced a partial weight phenotype that appeared robust upon switch to HFD (Fig. 4D). Consistent with published results (*1*), these animals displayed elevated BAT temperature, but normal food intake (Figs. 4E, F). Thus, *Rlim* loss specifically in GABAergic RIP-Cre^+^ neurons contribute to the observed increase in BAT thermogenesis (Fig. 2).

**Fig. 4:**
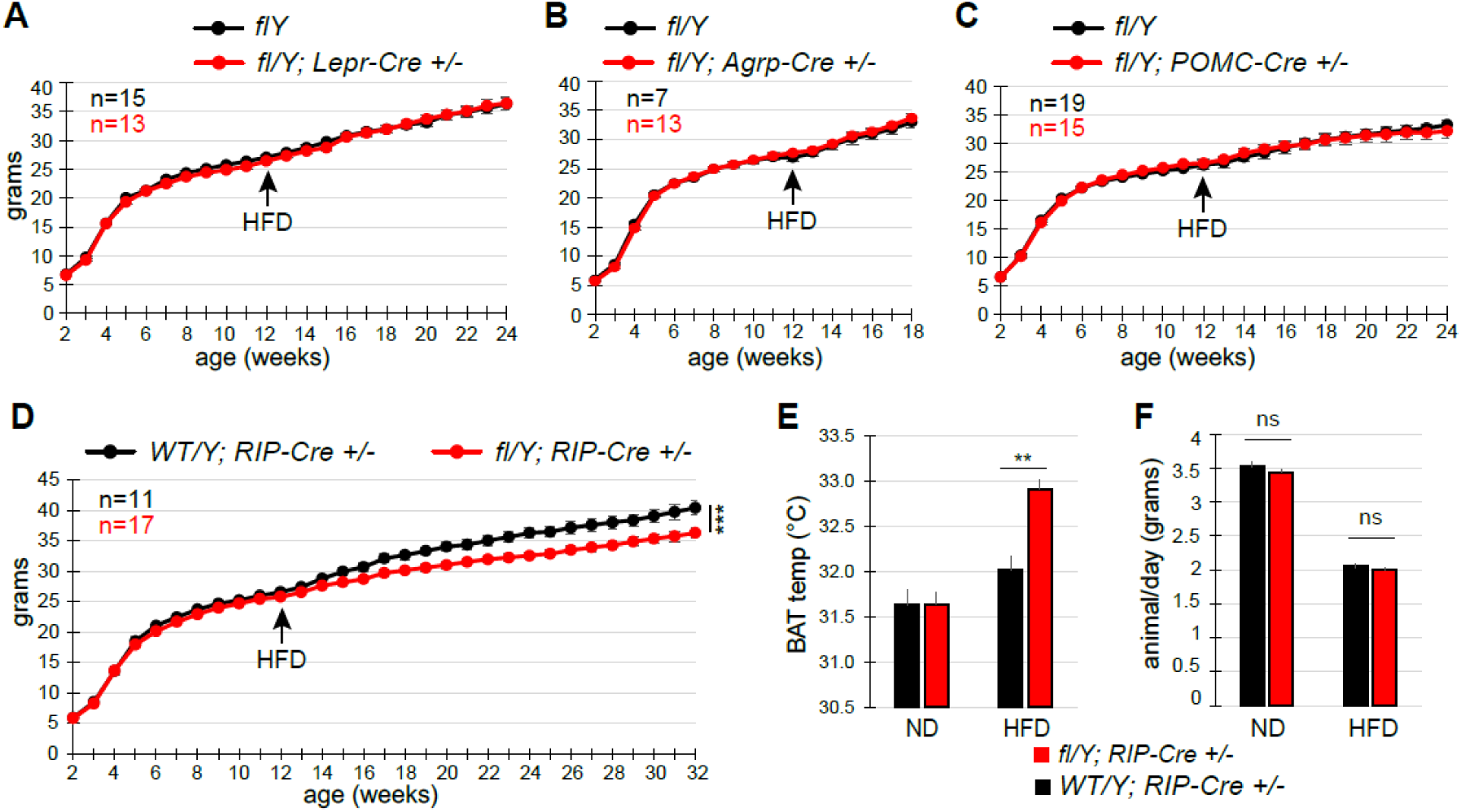
*Rlim* in GABAergic Rip-Cre^+^ neurons selectively regulates thermogenesis in BAT. Weight profiles: **Black =** fl/Y; **Red** = cKO/Y. P-values: Student’s t-test; *** = P<0.001; ** = P<0.01; Error bars indicate SEM. **A-C)** No effect on weights in animals with an induced *Rlim* cKO via LepR-Cre (A) AgRP-Cre (B) or POMC-Cre (C). **D)** Partial, HFD-dependent weight phenotype observed in animals with a RIP-Cre – induced *Rlim* cKO. fl/Y; RIP-Cre +/-were directly compared to control WT/Y; VIP-Cre +/-animals. **E, F)** Animals with a RIP-Cre mediated *Rlim* cKO fed HFD display increased BAT temperatures (E) but similar food intake when compared to controls (F).

### Rlim in RIP-Cre^+^ SCN neurons coordinates the diurnal rhythm of BAT thermogenesis

To identify RIP-Cre^+^ neurons in the mouse hypothalamus, where Cre expression is detected in several regions, including the Arc and the SCN (*1*), we generated *Rlim* fl/Y; EGFP^+/-^; RIP-Cre^+/-^ and control *Rlim* WT/Y; EGFP^+/-^; RIP-Cre^+/-^ animals. Validating these mice, immunohistochemistry (IHC) of hypothalamic brain sections in the SCN region confirmed broad RLIM protein expression including in RIP-Cre^+^ cells. The absence of RLIM in most EGFP^+^ neurons indicated efficient Cre-mediated deletion (Fig. 5A).

**Fig 5:**
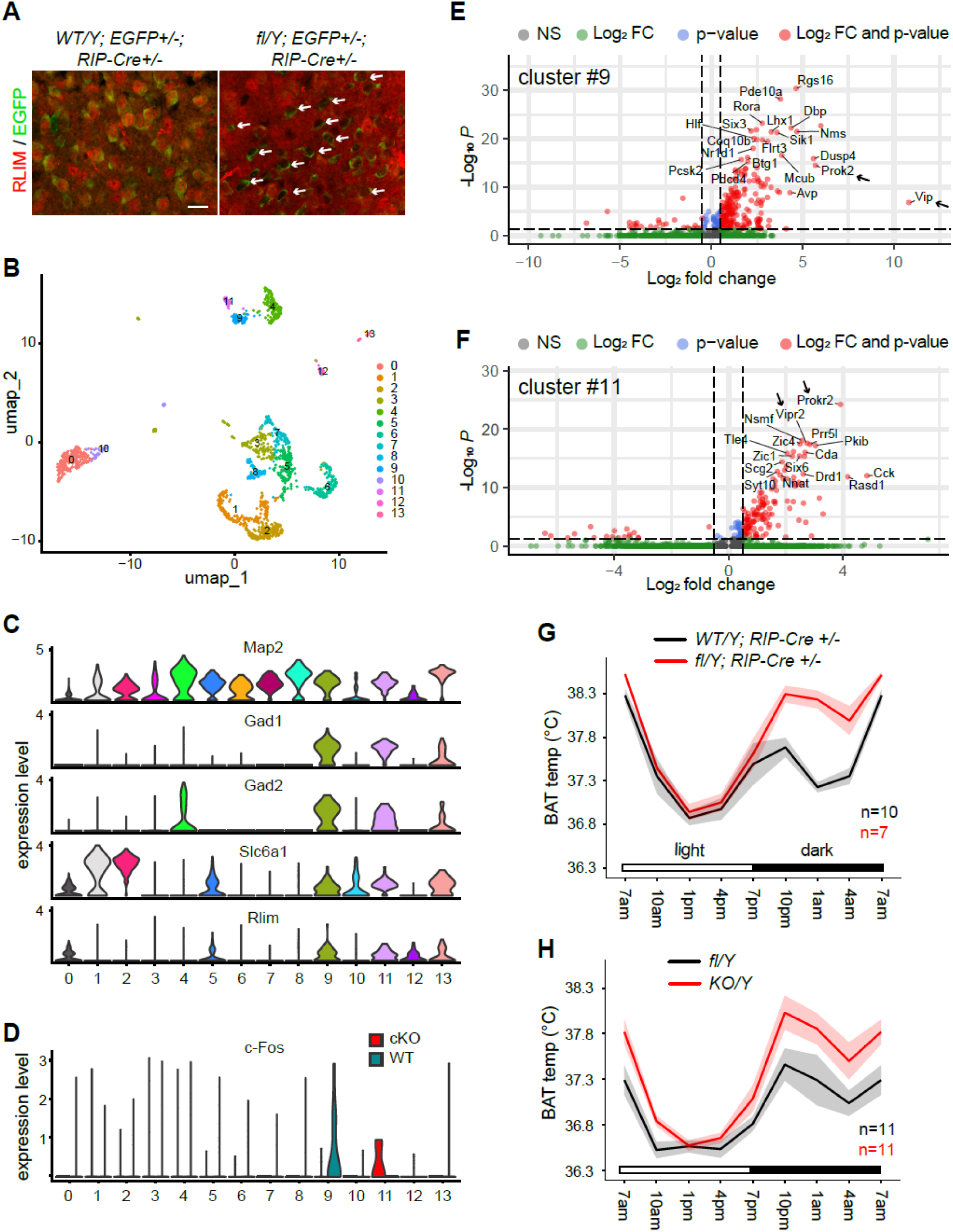
*Rlim* in RIP-Cre^+^ neurons of the SCN mediates diurnal regulation of thermogenesis. **A)** IHC using RLIM antibodies on hypothalamic sections in the SCN regions of *WT/Y; EGFP+/-; RIP-Cre +/-* and *fl/Y; EGFP+/-; RIP-Cre +/-* animals. Note EGFP+ cells with low/no RLIM staining (arrows) in the cKO, indicating correct targeting by the RIP-Cre transgene. Scale bar = 20 μm. **B)** UMAP projection of results (cKO and control) obtained by scRNA-seq across the entire dataset identifies 14 distinct cell clusters. **C)** Violin blots showing marker gene expression Map2 (neuronal), Gad1, Gad2, and Slc6a1 (GABAergic) identify clusters #9 and #11 among others, as containing GABAergic neurons, which express high relative levels of *Rlim* (cKO expresses 5.5kb out of 7.6kb total mRNA). **D)** Differential c-Fos expression between cKO and control with enriched c-Fos levels specifically in control cells of cluster #9 (P=2.66 × 10⁻⁹; likelihood ratio test (*33*)) and *Rlim* cKO neurons of cluster #11 (P<0.0003; likelihood ratio test (*33*)). **E, F)** Volcano blot comparing transcription profiles of cells in cluster #9 vs all other GABAergic neurons (E) or cells in cluster #11 vs all other GABAergic neurons (F). Only genes associated with the circadian clock are indicated. Arrows point at specific neuropeptides (E) and their receptors (F). **G)** BAT temperature profiles in mice with a RIP-Cre mediated cKO and RIP-Cre WT controls as determined by remote biotelemetry, measured in 3h intervals over a period of 72h. Note increased BAT temperature mostly during the dark phase (7PM-7AM). **H)** BAT temperature profiles in mice with a germline *Rlim* KO as determined by remote biotelemetry. Note similar BAT temperature patterns in G, H.

We performed scRNA-seq on these mice to elucidate the identity and location of RIP-Cre^+^ neurons in which *Rlim* is active, comparing primary hypothalamic EGFP^+^ neurons with and without *Rlim*. Hypothalamic brain regions of two 12 weeks-old mice per genotype, fed HFD were isolated via punches, cells dissociated in the presence of actinomycin D, followed by FACS isolation, yielding in around 1k EGFP^+^ cells. EGFP^+^ cells of each genotype were pooled and used for library preparation followed by sequencing. Analyses of sequencing results via the analysis package Seurat (*30*) separated 14 distinct cell clusters (#0-13; both in cKO and in WT) (Fig. 5B). A panel of 35 genes (curated from (*31*)), containing multiple markers for specific brain cell types was then used to unbiasedly annotate these clusters, identifying clusters containing GABAergic neuron subtypes (Fig. 5C). Because of a suspected persistent activity of *Rlim* in balancing energy homeostasis (Fig. S2), we compared WT and cKO neurons for changes in expression of the immediate early gene (IEG) c-Fos, a widely used marker of neuronal activity (*32*). Using a likelihood-ratio test for single cell feature expression (*33*), across the entire dataset we detected significant changes in c-Fos levels only in clusters #9 and #11 (Fig. 5D). Indeed, c-Fos levels were enriched in WT neurons of cluster #9 (P = 2.66 × 10⁻⁹) and in *Rlim* cKO neurons of cluster #11 (P<0.0003). Moreover, Gene Ontology (GO) Biological Pathway analysis of genes enriched in WT cells of cluster #9 (Fig. S5A) suggested altered neuronal activity and biophysical properties. To elucidate the identity of neuron subtypes involved, we carried out differential gene expression analyses across cells in cluster #9 or #11 vs all other GABAergic neurons. Surprisingly, these results revealed enriched expression of genes associated with the circadian clock in both clusters (Figs. 5E, F). Notably, neurons of cluster #9 expressed neuropeptides such as vasoactive intestinal peptide (VIP) and Prokineticin 2 (Prok2), while those in cluster #11 expressed their respective receptors (Vipr2; Prokr2) (*34–37*). GO biological process analysis of cluster #9 vs all other GABAergic neurons revealed enrichment of genes involved in rhythmic processes and circadian rhythm (Fig. S5B). Combined with the literature (*8*), these data place both clusters #9 and #11 containing neurons located in the SCN core, possibly participating in the circadian clock, and IHC confirmed RIP-Cre expression in a subset of neurons of the SCN including in the core region, where VIP is specifically expressed (Fig. S5C).

We used remote biotelemetry to investigate potential circadian effects mediated by *Rlim* on thermogenesis. Indeed, measuring BAT temperatures every three hours over a period of 3 days, we found robust increased BAT temperatures in HFD-fed mice with a RIP-Cre – induced *Rlim* cKO specifically at the dark (7PM – 7AM; active) phase but not during the light (7AM – 7PM; rest) phase (Fig. 5G). This pattern closely resembled that of germline *Rlim* KO/Y mice (Fig. 5H), indicating that RIP-Cre⁺ neurons mediate much of *Rlim*’s role in diurnal thermogenesis. Together, these results identify *Rlim* as a regulator of diurnal energy homeostasis in GABAergic RIP-Cre^+^ neurons.

### Rlim in VIP^+^ neurons of the SCN regulates the diurnal rhythm of food intake

Our scRNA-seq results suggested that *Rlim* in VIP^+^ neurons may influence energy homeostasis (Fig. 5E). In the hypothalamus, VIP is selectively expressed in neurons of the SCN core that represent the retinorecipient part of the circadian clock (*8, 38*). Indeed, VIP^+^ neurons are required for circadian rhythmicity (*39*), sending out GABAergic waves to various brain regions including the PVN (*40*), where signals regulating energy homeostasis are integrated (*5, 41*). Investigating *Rlim* function in VIP^+^ neurons, we directly compared *Rlim* WT/Y; VIP-Cre+/-with fl/Y; VIP-Cre +/-animals, thereby controlling for potential effects of this Cre transgene alone (*42*). *Rlim* fl/Y; VIP-Cre +/-mice exhibited a partial but robust HFD-dependent weight profile phenotype (Figs. 6A; S6A), with corresponding decreases in adipose tissue and liver mass (Figs. 6B; S6B, C). Examining BAT temperature profiles via remote telemetry, we found some effects of lack of *Rlim* on the daily pattern (Fig. S6D), but these changes did not recapitulate those observed in RIP-Cre cKO mice (Fig. 5G), suggesting additional involvement of other neuron subtype(s). To assess effects on feeding behavior, we measured food intake during the light (7AM – 7PM) and dark (7PM – 7AM) phases. Indeed, VIP-Cre – mediated *Rlim* cKO mice consumed significantly less food during the dark phase (Fig. 6C), mirroring the phenotype of germline *Rlim* KO mice (Fig. 6D). These findings indicate that *Rlim* in VIP⁺ SCN neurons regulates the diurnal rhythm of food intake and contributes to thermogenic control.

**Fig. 6:**
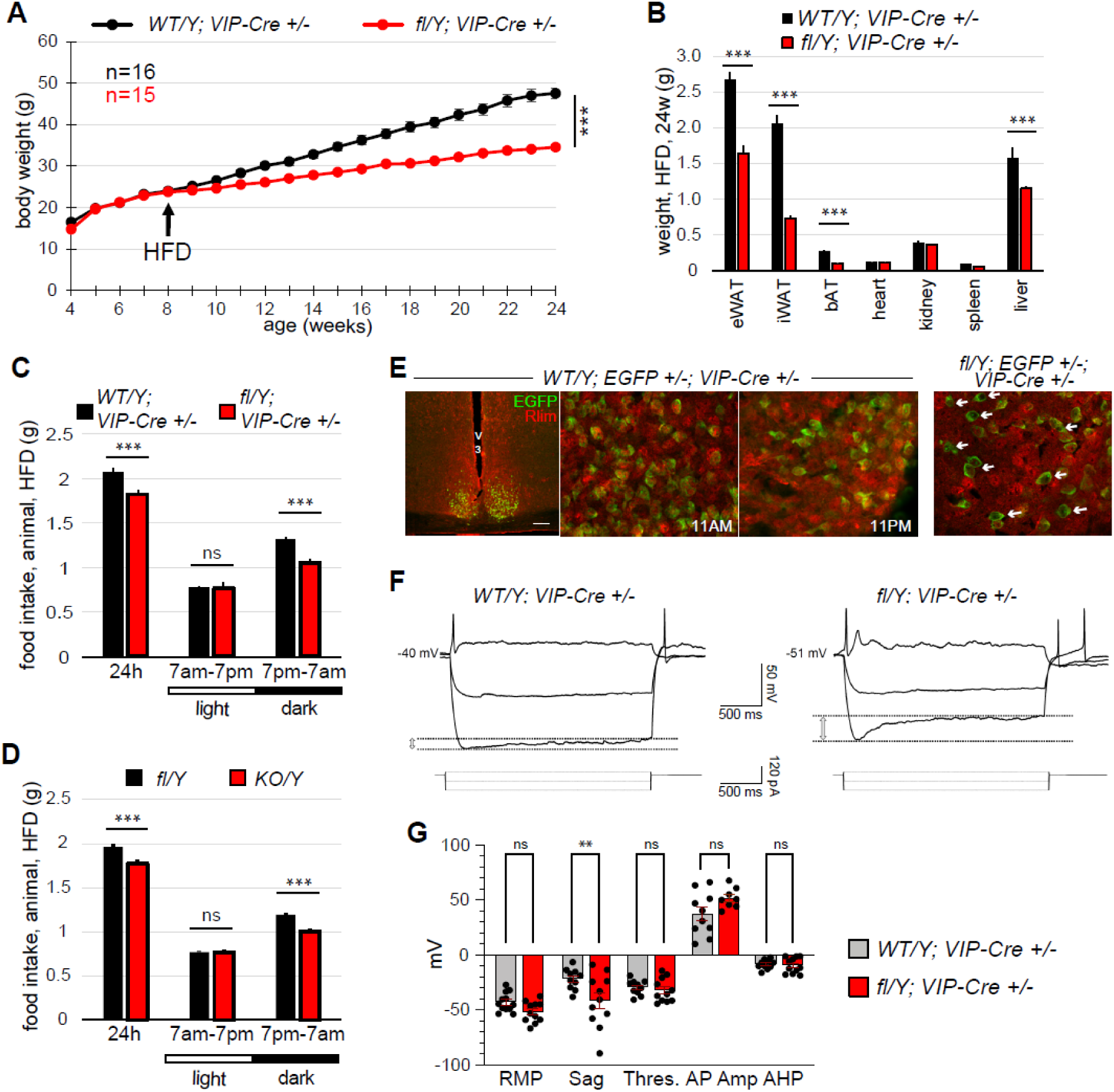
*Rlim* in VIP^+^ neurons of the SCN core mediates diurnal regulation of food intake. **A)** Body weight profiles comparing fl/Y; VIP-Cre +/-animals to WT/Y; VIP-Cre +/-controls. **B)** Comparative organ weights. Note significantly decreased weights of various adipose tissues as well as liver in cKO/Y animals. **C, D)** Food intake over the course of 24h in animals carrying a VIP-Cre – mediated *Rlim* cKO (C) or a germline KO (D) during the light and dark phase (7am-7pm and 7pm-7am, respectively). Note significantly less food intake of cKO and KO animals, specifically during the dark phase. *** = P<0.001; Student’s t-test. Error bars indicate SEM. **E)** IHC showing RLIM antibody staining and EGFP epifluorescence on SCN sections of *WT/Y; EGFP+/-; VIP-Cre +/-* and *fl/Y; EGFP+/-; VIP-Cre +/-* animals, collected and fixed at light and dark phases. Note no obvious difference in RLIM levels or cellular distribution during the light and dark phase in WT EGFP^+^ cells in the SCN core. Moreover, most EGFP^+^ cells in the cKO display low/no *Rlim* staining (arrows), indicating correct targeting by the VIP-Cre transgene. V3 = third ventricle; Scale bar = 100 μm. **F)** Representative voltage responses to negative and positive current steps in VIP neurons from control (WT/Y; VIP-Cre+/-, left panel) and cKO mice (fl/Y; VIP-Cre +/-, right panel) on sections prepared during the dark phase. Note the larger sag amplitude in the latter group. **G)** Average resting membrane potential (RMP; WT −42.92 ± 2.49 mV; n=12 cells; 3 mice and cKO −51.88 ± 2.85 mV; n=11 cells; 3 mice; adjusted p-value = 0.37; 95% CI [−22.64, 4.72]; One-way ANOVA Multiple Comparison Šidák), action potential threshold, (Thres, WT −29.43 ± 2.13 mV and cKO −32.12 ± 3.23 mV; adjusted p-value=0.99; 95% CI [−17.01, 11.63]; One-way ANOVA Multiple Comparison Šidák), action potential amplitude (AP Amp, WT; 37.26 ± 6.20 mV and cKO: 51.75 ± 3.27 mV; adjusted p-value=0.08; 95% CI [−1.06, 30.04] One-way ANOVA Multiple Comparison Šidák), after hyperpolarization (AHP, WT −8.48 ± 1.33 mV and cKO −9.42 ± 2.12 mV, adjusted p-value>0.99; 95% CI [−17.01, 11.63] One-way ANOVA Multiple Comparison Šidák), and sag (sag, WT −21.87 ± 3.01 mV and cKO −41.62± 7.32 mV; adjusted p-value = 0.002; 95% CI [−34.07, - 5.42] One-way ANOVA Multiple Comparison Šidák.

To further analyze functions of *Rlim* in VIP^+^ neurons, we generated *Rlim* fl/Y; EGFP^+/-^; VIP-Cre^+/-^ and control *Rlim* WT/Y; EGFP^+/-^; VIP-Cre^+/-^ mice. IHC confirmed EGFP expression specifically in hypothalamic SCN neurons and efficient *Rlim* deletion in the cKO group (Fig. 6E). Moreover, RLIM protein levels and subcellular localization did not obviously vary across light/dark cycles in control animals, suggesting that *Rlim* activity is not directly under circadian control.

Next, we used these animals to compare the biophysical properties of VIP EGFP^+^ neurons in the SCN with and without *Rlim* via whole-cell patch clamp electrophysiology. Due to the specific effects on food intake, these assays were performed on hypothalamic slices prepared during the dark phase from 3 independent mice per genotype kept in a reverse light room on a total of 12 WT and 11 cKO neurons. These experiments revealed similar resting membrane potentials of VIP EGFP^+^ neurons with or without *Rlim.* Injection of a positive (+20 pA) current pulse triggered action potentials (APs) that presented similar threshold, amplitude, and after hyperpolarization (AHP) (Figs. 6F, G). Moreover, the input resistance as well as the Rheobase were statistically similar between the two genotypes (Figs. S6E, F). As expected for SCN neurons during the dark phase (*43*), WT VIP⁺ neurons displayed low spontaneous firing rates. However, *Rlim*-deficient neurons showed a modest but statistically significant increase in firing frequency (Fig. S6G). Notably, the injection of a −100 pA current pulse initially hyperpolarized the membrane potential before inducing a sag whose amplitude was significantly larger in cKO (−41.62± 7.32 mV) compared to WT (−21.87 ± 3.01 mV) (Fig. 6G). This sag is mediated by hyperpolarization-activated cyclic nucleotide-gated (HCN) channels, which are critical for regulating neuronal excitability and rhythmic oscillatory activity (*44*). These findings indicate that *Rlim* modulates the intrinsic biophysical properties of VIP⁺ SCN neurons, potentially influencing their role in circadian regulation of energy homeostasis.

## Discussion

Using a genetic approach, our study reveals a central role for the E3 ubiquitin ligase *Rlim* in regulating diurnal rhythms of energy homeostasis in mice. While previous *in vivo* studies have highlighted sex-specific functions of *Rlim* in various physiological processes (*18, 22, 25*), our findings demonstrate that its influence on weight gain is comparable in both sexes. This suggests a broader, sex-independent role for *Rlim* in maintaining energy balance.

We provide genetic evidence that the effects of *Rlim* deletion on weight gain are at least partially dependent on its sustained expression (Fig. S2). This is supported by the absence of significant differences in VIP-Cre or RIP-Cre cell numbers in the SCN between cKO and control animals, alongside notable alterations in the biophysical properties and activity of VIP neurons lacking *Rlim* (Fig. 6). However, while these findings are consistent, we cannot fully exclude minor developmental contributions to the observed phenotype.

Our data show that *Rlim* is essential for the daily regulation of thermogenesis and food intake in specific neuronal subtypes, particularly under high-fat diet (HFD) conditions. Although HFD alone can influence peripheral circadian oscillators, it does not substantially disrupt the central circadian clock (*45*). However, our results indicate that the combination of HFD and *Rlim* deficiency significantly alters circadian processes, including diurnal energy homeostasis. These findings warrant further investigation into *Rlim*’s potential roles in other circadian-controlled behaviors, such as physical activity and/or sleep. Importantly, we also identify HFD-independent functions of *Rlim* in GABAergic neurons. This is supported by weight gain phenotypes in cKO mice driven by Sox2-Cre, Nestin-Cre, and Vgat-Cre, as well as in germline KO animals (Figs. 1A; 3A, B; S1), suggesting that *Rlim* may act in additional neuronal subtypes beyond those currently characterized.

Concerning the HFD-dependent regulation of energy homeostasis by *Rlim*, diurnal regulation of thermogenesis but not food intake appears mostly mediated by RIP-Cre^+^ neurons located in the SCN, as germline *Rlim* KO mice exhibit a similar BAT temperature pattern when compared to RIP-Cre-mediated cKO mice (Figs. 5G, H). This finding is consistent with published data (*1*) and supports a broader role for hypothalamic RIP-Cre^+^ neurons, including those in the arcuate nucleus and SCN, in thermogenic control. Notably, VIP-Cre^+^ neurons lacking *Rlim* do not replicate the thermogenic phenotype observed in RIP-Cre cKOs (Fig. S6D), implicating multiple RIP-Cre^+^ neuronal subtypes—such as those in clusters #9 and #11—in this regulation. Indeed, neurons in these clusters are likely interconnected (*8*), further supporting their coordinated role. However, contributions from other neuronal populations in the regulation of BAT temperature cannot be excluded.

Regarding the daily rhythm of food intake, our data highlights a critical role for *Rlim* in VIP^+^ neurons of the SCN core. Consistent with such a role, these neurons send out GABAergic waves to various brain regions including the PVN (*40*), where signals regulating energy homeostasis are integrated (*5, 41*). Indeed, we provide evidence that the *Rlim* deletion alters both the transcriptomic and biophysical properties of these neurons. Specifically, we observed significant changes in mRNA levels of ion channel genes in cluster #9, including downregulation of HCN1 and HCN2, thereby possibly explaining the reduced sag observed in VIP^+^ neurons lacking *Rlim* (Figs. 6F, G). Given the role of *I*h currents—mediated by HCN channels—in neuronal oscillations (*43, 44*), these findings suggest a mechanistic link between *Rlim* and the rhythmic regulation of energy balance.

In conclusion, our findings establish *Rlim* as a key epigenetic regulator of diurnal energy homeostasis, acting through distinct GABAergic neuronal populations in the SCN. By modulating thermogenesis and food intake via RIP-Cre^+^ and VIP-Cre^+^ neurons, respectively, *Rlim* emerges as a critical node linking circadian timing to metabolic control.

## Material and Methods

### Mice and generation of animals

Mice were bred and generally maintained in a C57BL/6 background, whenever possible. *Rlim* germline KO, *Rlim fl/fl*, and mice with a Sox2-Cre – mediated *Rlim* cKO have been described (*17, 19, 20*). Transgenic mouse lines purchased from The Jackson laboratory were: *Sox2-Cre* (JAX #008454), *GFAP-Cre* (JAX #024098), *Sf1-Cre* (JAX #012462); *Ubc-CreERT2* (JAX #007001); *Nestin-Cre* (JAX #003771); *adiponectin-Cre* (JAX # 010803); *POMC-Cre* (JAX #005965); *CamK2-Cre* (JAX #005359); *AgRP-Cre* (JAX #012899); *Chat-Cre* (JAX #006410); *DAT-Cre* (JAX# 006660); *Vgat-Cre* (JAX #016962); *LepR-Cre* (JAX #008320); *Ins1-Cre* (JAX #0026801); *Vil1-Cre* (JAX #021504); *Sox10-Cre* (JAX #025807); *RIP-Cre* (JAX #003573); *Pdx1-Cre* (JAX #014647); *VIP-Cre* (JAX #031628); *fl-Stop-fl EGFP* (JAX #024750). *Sim1-Cre* mice (kind gift of B. Lowell) have been described (*27*). A list of mouse lines used in this study is provided in Table S1. Cre-mediated *Rlim* cKO animals were generated following several strategies: To generate hemizygous *Rlim* cKO males, sires heterozygous for the Cre transgene and WT for *Rlim* (*Rlim* WT/Y; Cre+/-as obtained from JAX) were mated with *Rlim* fl/fl dames. The resulting F1 male littermates (fl/Y; Cre+/- and fl/Y) were directly compared, and potential Cre-only effects were then tested in a WT/Y background. Homozygous RIP-Cre+/+ or VIP-Cre+/+ males as obtained from JAX were mated to fl/fl or WT/WT females and effects in F1 male fl/Y; Cre+/-offspring were directly compared to F1 male WT/Y; Cre+/-. Female mice systemically lacking *Rlim* via Sox2-Cre were generated by targeting the maternally transmitted fl allele via a paternally transmitted Sox2-Cre in epiblast tissues after imprinted iXCI as described (*19, 20*). Systemic Cre expression in adult *Rlim fl/Y*; *Ubc-CreERT2* males was induced by tamoxifen injections as recommended by the Jackson Laboratory (https://www.jax.org/research-and-faculty/resources/cre-repository/tamoxifen). Mice were housed in the animal facility of UMass Chan Medical School in a room set at ∼21°C with food and water available ad libitum under a daily 12h light/dark cycle (7am-7pm / 7pm-7am). Animals used for electrophysiological studies were kept in a reverse light room (7pm-7am / 7am-7pm). All mice were utilized according to NIH guidelines and those established by the UMMS Institute of Animal Care and Usage Committee (IACUC).

### In vivo experiments

Mice were fed normal a chow diet (ND) or high-fat-diet (HFD), fat Calories 60%, Bio-Serv, S3282, as indicated. For elucidating the weight profiles, animals were weighed on a weekly basis. Scapular BAT temperatures were determined using an infrared thermal camera (FLIR T420) on lightly anesthetized mice. Images were analyzed with FLIR tools. BAT temperature measurements via remote telemetry were carried out using a system from Unified Information Devices, Inc. (reader URH-1HP; microchips UCT-2112-24) according to the manufacturer’s instructions. Measurements were recorded every 3h over a period of 72h. To minimize disturbance of the circadian cycle of animals, nighttime measurements were carried out under red light. Studies involving metabolic cages and 1H-MRS were performed by the Mouse Metabolic Phenotyping Center at the University of Massachusetts Chan Medical School. The mice were housed in metabolic cages (TSE Systems) under controlled conditions including temperature and lighting. Energy expenditure, respiratory exchange ratio, and physical activity were determined. The body composition (lean vs fat mass) of live mice was determined using 1H-MRS (Echo Medical System). The analysis of lipids in mouse feces was performed as described (*46*).

### Antibodies, Immunohistochemistry and histology on tissue sections

Immunohistochemical staining of paraffin-embedded tissue sections (liver, BAT) was carried out as previously reported (*18*). IHC on mouse brain sections was performed essentially as described (*47*). Briefly, mice were transcardially perfused with 10% formalin, brains were removed and postfixed for 1 d at 4°C, followed by dehydration in PBS/30% sucrose at 4°C for 2–3 d. Then, 30 μm coronal sections were taken using a sliding microtome (Leica SM2010 R) and slices were blocked in PBS with 0.2% Triton X-100, 5% normal goat serum. Images were acquired on a Nikon confocal microscope (Nikon, ECLIPSE Ti2) and Leica Thunder. Primary antibodies used for immunostainings were rabbit RLIM (*11, 12*), UCP1 (Sigma, CU6382), c-Fos (Synaptic Systems, 226003), and VIP (Invitrogen, PA578224). Secondary antibodies were Alexa Fluor® 488 Donkey Anti-Rabbit IgG (Invitrogen, A21206) and Alexa Fluor® 568 Goat Anti-Rabbit IgG (Invitrogen, A11036). H&E staining of paraffin-embedded tissue sections was performed as previously described (*18*). Staining of liver sections using Oil red O (Sigma, 1024190250) was carried out according to manufacturers’ instructions.

### scRNA-seq

scRNA-seq experiments were performed using *Rlim* fl/Y; EGFP^+/-^; RIP-Cre^+/-^ and control *Rlim* WT/Y; EGFP^+/-^; RIP-Cre^+/-^ animals. To isolate single cells for scRNA-sequencing, punches of hypothalamic brain regions (approx. 4 mm^3^) of two mice per genotype (12 weeks-old, HFD, non-fasted, housed at RT, brains harvested at 10:30 AM) were collected in DPBS supplemented with 1g/L D-glucose, 36 mg/L sodium pyruvate (Gibco) and 45 μM actinomycin-D (Sigma). To generate a single cell suspension, dissociation was performed as previously reported (*48*) on the Singulator 100 (S2 Genomics) following the manufacturer’s recommended protocol for mouse brain dissociation using the mouse brain enzyme reagent (S2 Genomics) resuspended in DMEM (Gibco). Briefly, the samples were rinsed in DPBS supplemented with actinomycin-D, loaded into a cell isolation cartridge (S2 Genomics), and run on the instrument ensuring that the DMEM buffer for collection was supplemented with actinomycin-D. The cell suspension was recovered from the cartridge and centrifuged at 300g for 10 minutes at 4 °C and the supernatant was carefully discarded. Debris removal and clean up was performed as previously described (*49*) and after centrifugation, the cell pellet was resuspended in DPBS supplemented with 0.04% BSA and actinomycin-D. The single cell suspension was filtered through a 30 μm MACS SmartStrainer (Miltenyi Biotec) cell strainer and the cells were counted in a Countess II FL (Invitrogen) using Trypan blue stain (Invitrogen). Fluorescence-activated cell sorting of the samples was done in the UMASS Chan Flow core facility using a BD FACSFusion Cell Sorter, isolating EGFP^+^ cells (>95% efficiency). The eluants were prepared for Gel Beads-in-emulsion (GEMs) (10x Genomics) generation and barcoding.

The Chromium Next GEM Single Cell 3’ v3.1 kit (10x Genomics) was used to generate Genomic libraries for scRNA-seq following the manufacturer’s protocol. Briefly, the barcoded Single Cell 3’ v3.1 gel beads were combined with the single cell suspension in the master mix and partitioning oil and the GEM generated by the Chromium Controller (10X Genomics). The GEMs were transferred from the chip into a tube strip and then reverse transcribed in a thermal cycler to generate cDNA. Post GEM-RT cleanup was performed, and the cDNA was amplified. After cDNA clean-up, 1 μL of the sample was run on a Bioanalyzer High Sensitivity chip (Agilent) for quality control and quantification. The cDNA was used to generate 3’ gene expression libraries using the Chromium Next GEM Single Cell 3’ Library kit v3.1 (10x Genomics), which were then shipped to Novogene Inc. for sequencing after post library construction quality control on an Agilent Bioanalyzer (Agilent).

For data analyses, Cell Ranger (10x Genomics) was used to aggregate biological replicates, perform alignment, filter, count barcodes and Unique Molecular Identifiers (UMIs). Neuronal identity was confirmed by observing canonical neuronal marker genes (Map2, NeuN and Thy1) across the dataset. Clustering, differential gene expression analysis, and integrative analysis of wildtype and knockout conditions were performed using the Seurat V4 package (*28*). Low quality cells (genes < 200 or genes > 7000, or percentage of mitochondrial genes > 20%) were removed prior to analysis. 1,774 cells across wildtype and *Rlim* cKO samples were then used for downstream analysis (*28, 29*). Briefly, gene counts were scaled by the cellular sequencing depth (total UMI) with a constant scale factor (10,000) and then natural log transformed (log1p). 2,000 highly variable genes were selected in each sample based on a variance stabilizing transformation. Anchors between individual data were identified and correction vectors were calculated to generate an integrated expression matrix, which was used for subsequent clustering. Integrated expression matrices were scaled and centered followed by principal component analysis (PCA) for dimensional reduction. PC1 to PC18 were used to construct nearest neighbor graphs in the PCA space followed by Louvain clustering to identify clusters (resolution = 0.5). For visualization of clusters, Uniform Manifold Approximation and Projection (UMAP) was generated using the same PC1 to PC18. Differentially expressed genes between genotypes were determined using a likelihood-ratio test for single cell feature expression (*31*) (see Data S1, S2).

### Electrophysiology

Coronal slices were prepared from fresh brain tissue from 12-week-old male mice at Zeitgeber time (ZT) 3. Following intracardiac perfusion with an ice-cold N-methyl-D-glucamine-based (NMDG) solution (see below), we rapidly removed and transferred the brain to a cold (∼ +0.5°C) oxygenated (95% O_2_ and 5% CO_2_) cutting solution of the following composition (in mM): 95 N-methyl-D-glucamine (NMDG), 2.5 KCl, 1.25 NaH_2_PO_4_ · H_2_O, 30 NaHCO_3_, 20 HEPES, 25 D-Glucose, 2 thiourea, 5 Na+ -ascorbate, 3 Na+ - pyruvate, 0.5 CaCl_2_, 10 MgSO_4_ · 7H_2_O. Coronal sections containing the suprachiasmatic nucleus (SCN) were sliced into 200 μm sections with a Vibroslicer (VT1200, Leica MicroInstruments; Germany). Slices were immediately transferred to an incubation chamber and left to recuperate in the NMDG-based solution for 20 min at 30°C before being moved into a chamber containing oxygenated artificial cerebrospinal fluid (aCSF; in mM): 126 NaCl, 2.5 KCl, 1.3 NaH_2_PO_4_· H2O, 1 MgCl_2_, 2 CaCl_2_, 26 NaHCO_3_, 10 D-Glucose, at room temperature. Slices were left in this chamber for at least one hour before being placed in a recording chamber and perfused with oxygenated aCSF at a constant rate of 2–3 ml/min at room temperature (∼21°C). We visualized neurons in infrared differential interference contrast (60×, IR-DIC) video microscopy using a fully motorized upright microscope (Scientifica; England).

Concerning electrophysiology, whole-cell patch clamp recordings of VIP EGFP+ neurons were performed in a recording chamber perfused with oxygenated (95% O_2_ and 5% CO_2_) artificial cerebral spinal fluid (aCSF) with the following composition (mM): 126 NaCl, 2.5 KCl, 1.3 NaH_2_PO_4_ · H_2_O, 1 MgCl_2_, 2 CaCl_2_, 26 NaHCO_3_, 10 D-Glucose, at a rate of 2-3 ml/min at room temperature. We identified EGFP-labeled VIP neurons in the SCN core using a broad-spectrum blue light (425-500 nm; 450 nm peak) source (pE- 300white, CooLED, NY, United States) passed through a 535/50 emission filter and a 500/20 excitation filter (Chroma, Bellows Falls, VT) at 60X magnification. Briefly, borosilicate glass electrodes (1.5 mm OD, 7– 14 MΩ resistance) were filled with an internal solution containing (mM): 120 K-methanesulfonate; 20 KCl; 10 HEPES; 2 ATP, 1 GTP, and 12 phosphocreatine. Following seal rupture, series resistance was 25.2 ± 1.1 MΩ, fully compensated in current clamp recording mode, and periodically monitored throughout recording sessions. Recordings with changes of series resistance larger than 20% were rejected. Current-Voltage traces were recorded with the following parameters: 14 sweeps with an increasing injected current of +20 pA with an initial injected current of −100 pA. Cellular biophysical properties (i.e. action potentials, rheobase, input resistance, and refractory period) were measured by holding the membrane potential at the cellular resting membrane potential. Voltage and current traces in whole- cell patch-clamp were acquired with an EPC10 amplifier (HEKA Elektronik; Germany). Sampling was performed at 10 kHz and digitally filtered voltage and current traces were acquired with PatchMaster 2.15 (HEKA Elektronik; Germany) at 2 kHz. All traces were subsequently analyzed offline with FitMaster 2.15 (HEKA Electronik; Germany). The experimenter was blinded during patch clamp experiment and analysis but was unblinded during data interpretation.

## Author contributions

Conceptualization: I.B. and F.W.; Methodology: F.W. and M.Z., Investigation, F.W., P.M., M.Z. P.M., T.L., H.L. and D.R.W.; Supervision: I.B., T.T., G.E.M., D.A.G. and J.K.K.; Writing: I.B., F.W.

## Acknowledgments

We are grateful to C. Anaclet, R. Davis, P. Emory, L. Ferrari, S.M. Jung, H. Li, J. Mao, V. Navarro, A. Tapper, Y.-X. Wang, and D. Weaver for advice, methodological help and/or discussion. *Sim1-Cre* transgenic mice were a kind gift of B. Lowell. I.B., D.A.G and J.K.K. are members of the University of Massachusetts DERC (DK32520). Part of this study was performed at the National Mouse Metabolic Phenotyping Center (MMPC) at the University of Massachusetts Chan Medical School, funded by NIH grant 5U2C-DK093000. This work was supported by NIH grants R35GM145263 to I.B., R01DK094004 and R01DK127175 to D.A.G., and R01DK133772 to J.K.K.

## Supplementary Figures

**Fig. S1:**
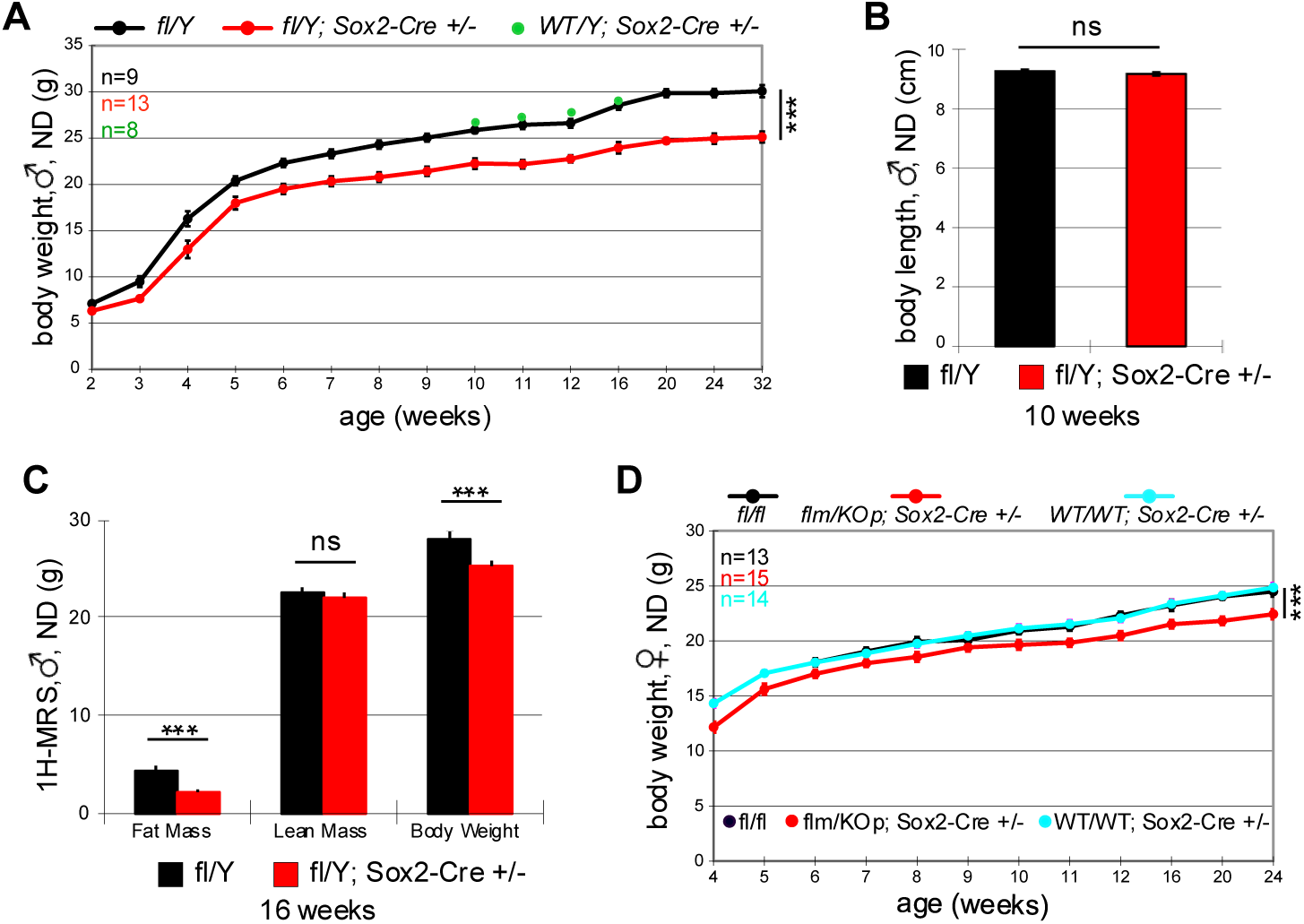
Male and female mice lacking *Rlim* are lean. *** = P<0.001; Student’s t-test. Error bars indicate SEM. **A)** Profiles of body weights of male mice with a Sox2-Cre – mediated systemic *Rlim* cKO in comparison with fl/Y and and Sox2-Cre-only control males. **B)** Similar body length between cKO and control animals. **C)** Significantly less fat mass but not lean mass in males lacking *Rlim* as revealed by 1H-MRS. **D)** Profiles of body weights of female mice systemically lacking *Rlim*.

**Fig. S2:**
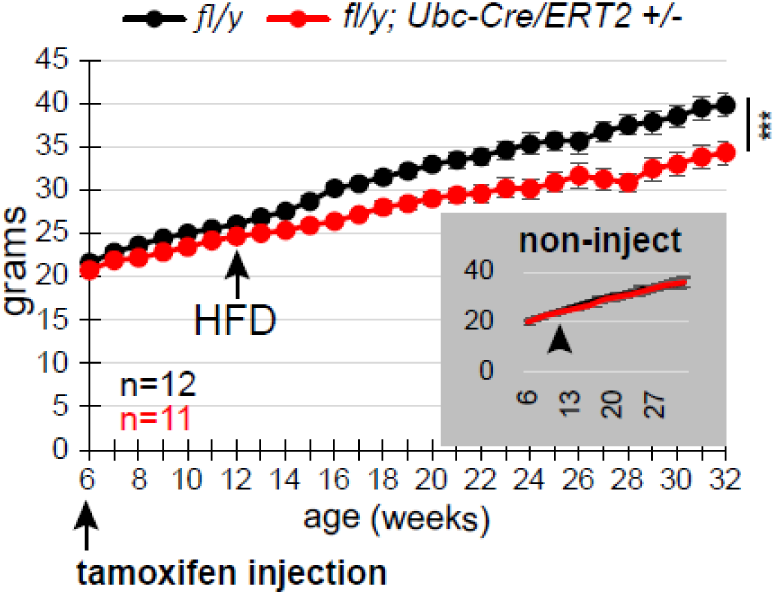
A persistent activity of *Rlim* is involved in the balancing of energy homeostasis. Weight profiles of Ubc-Cre/ERT2 fl/Y males and fl/Y-only control littermates, with a systemic *Rlim* cKO induced in adults via tamoxifen-injections. Non-injected animals served as control (inset). Results indicate that effects in *Rlim* KO mice are at least partially based on the disturbance of a persistent activity. *** = P<0.001; Student’s t-test. Error bars indicate SEM.

**Fig. S3.**
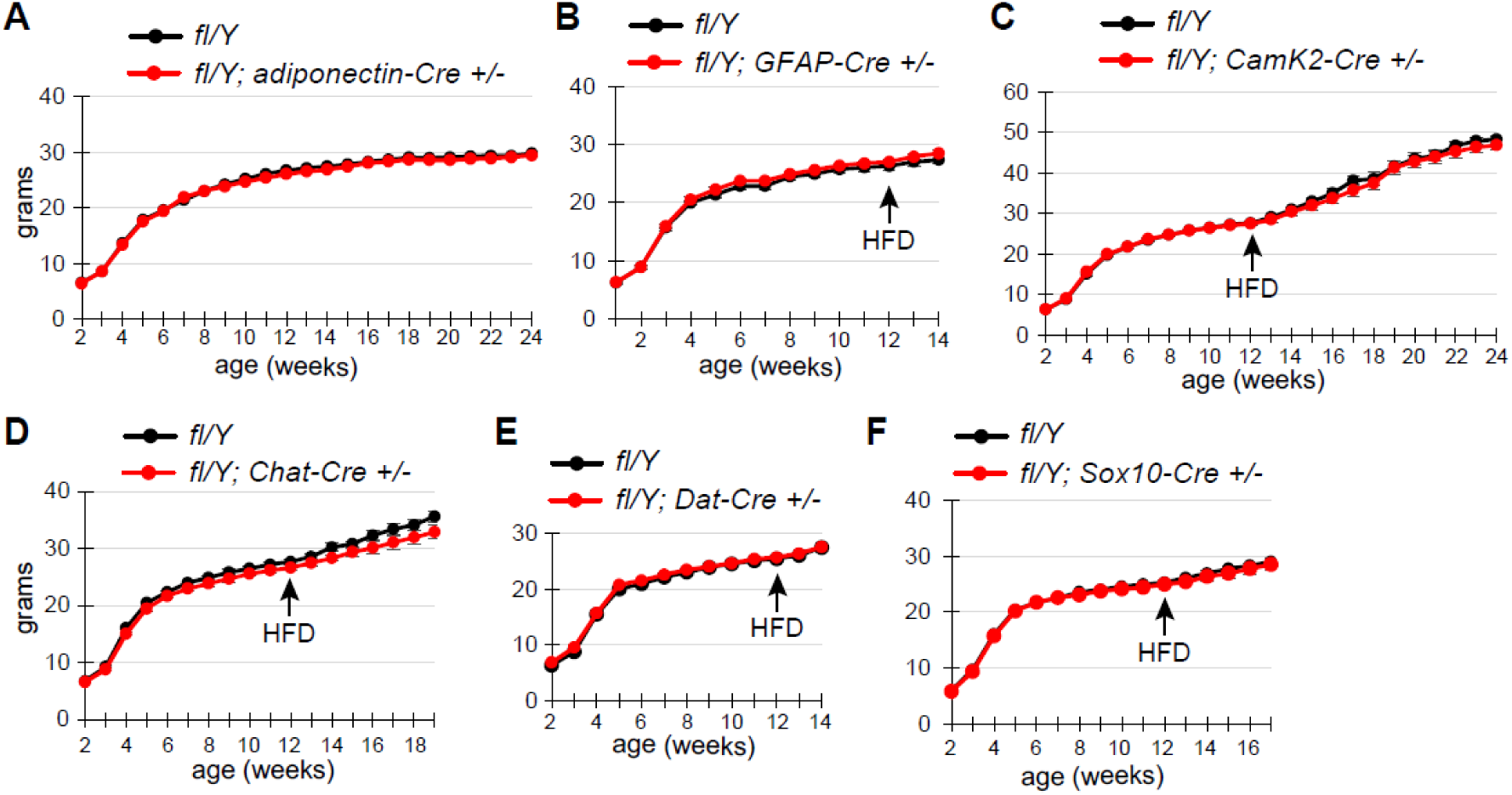
*Rlim* acts in GABAergic neurons to regulate energy homeostasis. Weight profiles: **Black =** fl/Y; **Red** = cKO/Y). *** = P<0.001; ** = P<0.01; Student’s t-test; error bars indicate SEM. Targeting of the *Rlim* cKO in: **A)** White and brown adipocytes via adiponectin-Cre (adipoq-Cre), **B)** Astrocytes via GFAP-Cre, **C)** Excitatory neurons of many brain regions via CamK2-Cre, **D)** Cholinergic neurons via Chat-Cre **E)** Dopaminergic neurons via DAT-Cre, and **F)** Neural crest cells including cells of the enteric nervous system via Sox10-Cre.

**Fig. S4.**
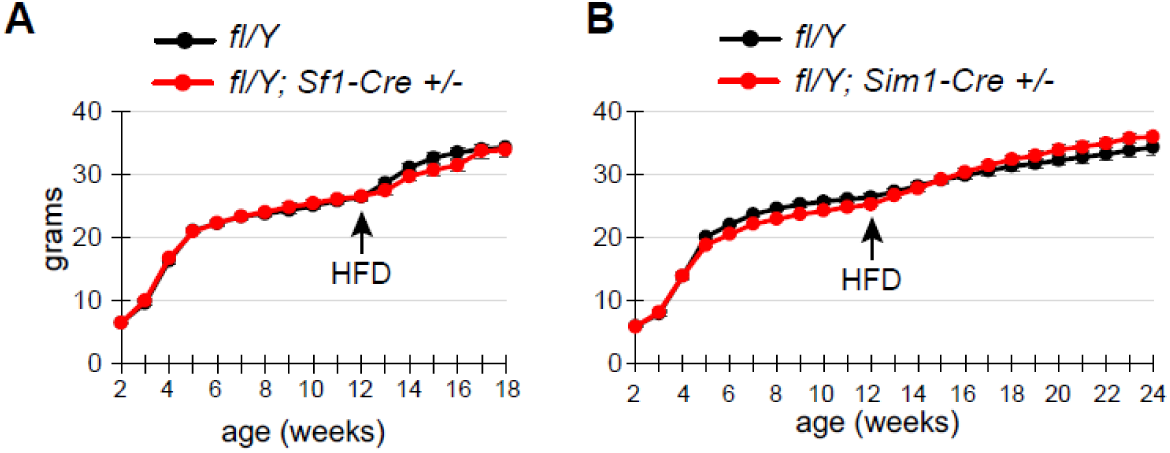
*Rlim* in RIP-Cre^+^ neurons regulates BAT thermogenesis. Weight profiles: **Black =** fl/Y; **Red** = cKO/Y). *** = P<0.001; ** = P<0.01; Student’s t-test; error bars indicate SEM. Targeting of the *Rlim* cKO in: **A)** Neurons in the hypothalamic VMN region via Sf1-Cre, and **B)** Neurons in the hypothalamic PVN region via Sim1-Cre.

**Fig. S5:**
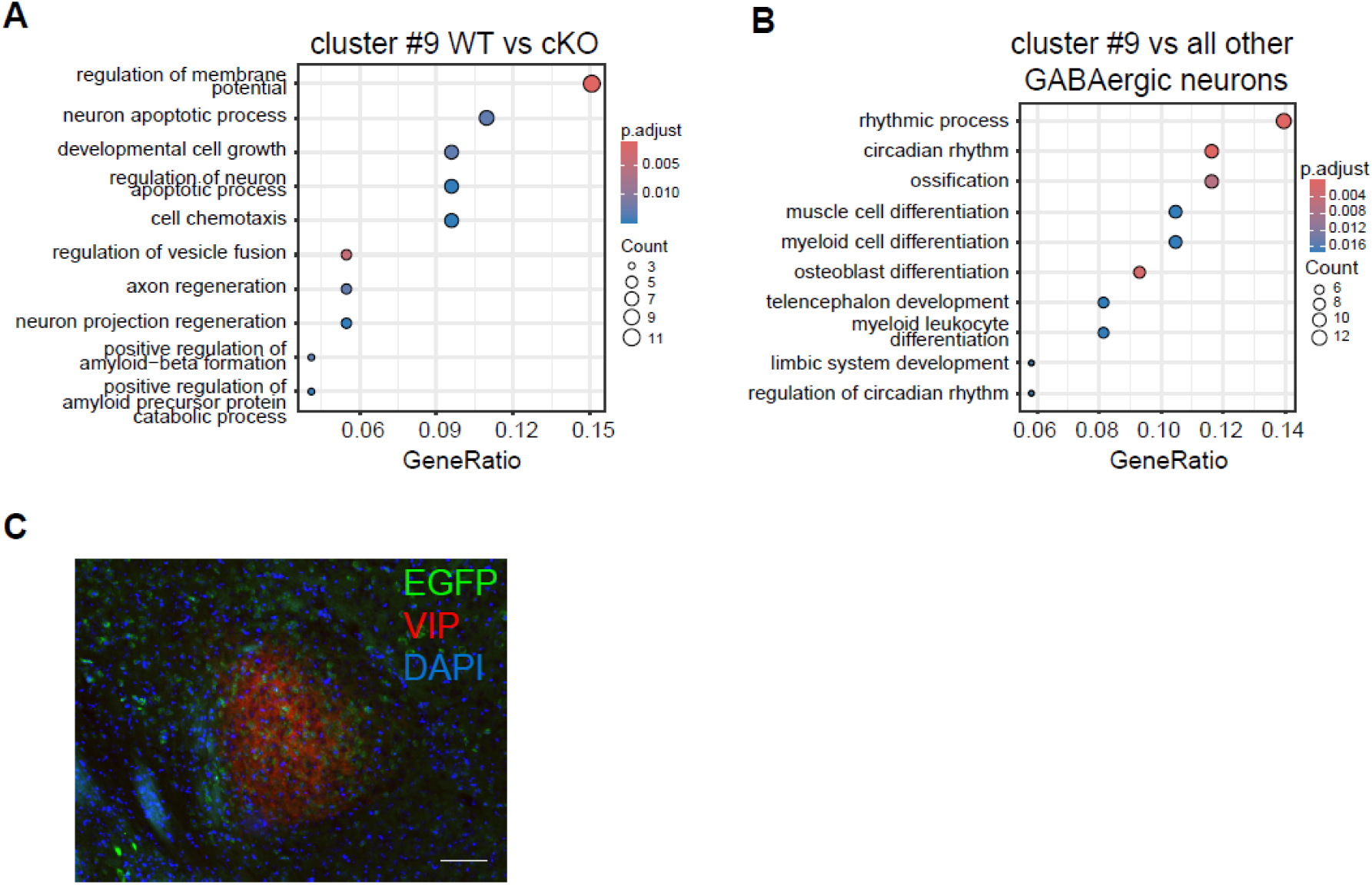
*Rlim* acts in SCN neurons. **A)** Biological pathway Gene Ontology terms of differential transcription profiles of genes enriched in RIP-Cre WT controls vs RIP-Cre mediated *Rlim* cKO/Y of cluster #9 as determined via scRNA- seq. **B)** Biological pathway Gene Ontology terms of transcription profiles comparing cells of cluster #9 vs all other GABAergic neurons. **C)** IHC using VIP antibodies on hypothalamic sections of the SCN regions of *WT/Y; EGFP+/-; RIP-Cre +/-* animals. Note EGFP^+^ cells in regions of VIP expression, indicating a RIP-Cre^+^ cell population within the SCN core. Scale bar = 60 μm.

**Fig. S6:**
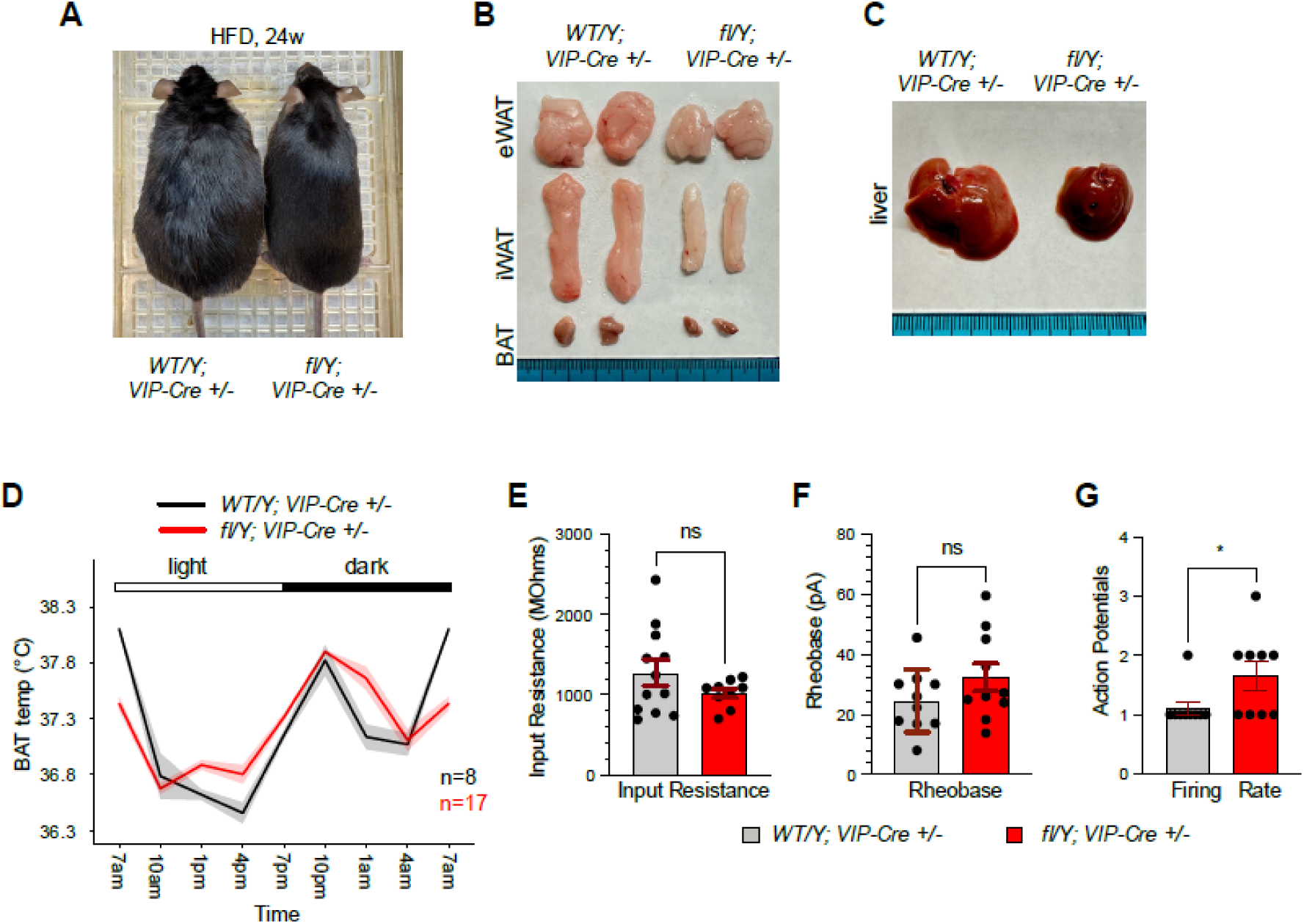
*Rlim* in SCN VIP+ neurons mediate diurnal regulation of food intake. WT/Y; VIP-Cre +/- control and fl/Y; VIP-Cre +/- males were fed 8 weeks with ND and then switched over to HFD. **A)** Appearance of WT/Y; VIP-Cre +/- and fl/Y; VIP-Cre +/- animals at 24 weeks (HFD). **B)** Representative adipose tissues isolated from animals at 24w. **C)** Representative livers of animals at 24w. **D)** BAT temperature profiles in mice with a VIP-Cre mediated *Rlim* cKO and VIP-Cre *Rlim* WT controls as determined by remote biotelemetry, measured in 3h intervals over a period of 72h. **E)** Average VIP neurons input resistance in WT (1272 ± 156 Mohms) and cKO (1017 ± 57 Mohms) mice (p-value=0.19; t=1.35, df=19; two-tailed unpaired t-test). **F)** Average VIP neurons rheobase in WT (24.57 ± 3.32 pA) and cKO (32.42± 4.64 pA) mice (p- value=0.19; t=1.38, df=18; two-tailed unpaired t-test). **G)** Marginally but significantly higher firing rate as counted by the number of action potentials in VIP+ neurons lacking *Rlim* (p-value= 0.048; t=2.13, df=16; two-tailed unpaired t-test).

**Table S1:** Mouse lines.

| # | Cre | source;<br>JAX # | regions affected | weight<br>phenotype |
| --- | --- | --- | --- | --- |
| 1 | Adipoq | 010803 | all adipose tissues | no |
| 2 | AGRP | 012899 | Arc neurons, GABAergic | no |
| 3 | CamK2 | 005359 | brain, excitatory neurons | no |
| 4 | Chat | 006410 | cholinergic neurons, ENS | no |
| 5 | DAT | 006660 | dopaminergic neurons | no |
| 6 | GFAP | 024098 | astrocytes | no |
| 7 | Ins1 | 026801 | pancreatic $\beta$ cells | no |
| 8 | LepR | 008320 | leptin receptor+ cells | no |
| 9 | Nestin | 003771 | neural, other tissues | strong |
| 10 | Pdx1 | 014647 | $\beta$ cells, brain regions | no |
| 11 | POMC | 005965 | Arc neurons, ant. pituitary | no |
| 12 | RIP | 003573 | $\beta$ cells, hypothalamus | medium |
| 13 | Sf1 | 012462 | VMN, pituitary, adrenal | no |
| 14 | Sim1 | Ref. 27 | PVN neurons | no |
| 15 | Sox2 | 008454 | ubiquitous | strong |
| 16 | Sox10 | 025807 | neural crest, ENS | no |
| 17 | Ubc-ERT2 | 007001 | ubiquitous | medium |
| 18 | Vgat | 016962 | GABAergic, ENS, EE, b cells | strong |
| 19 | Vil1 | 021504 | gut epithelial cells incl. EE | no |
| 20 | VIP | 031628 | SCN, brain, pancreas, gut | medium |
| <u>other</u> |  |  |  |  |
| 21 | Rlim fl/fl | Ref. 17 |  |  |
| 22 | EGFP fl-stop-fl | 005667 |  |  |

## References

1. D. Kong et al., GABAergic RIP-Cre neurons in the arcuate nucleus selectively regulate energy expenditure. Cell 151, 645–657 (2012).

2. Y. Aponte, D. Atasoy, S. M. Sternson, AGRP neurons are sufficient to orchestrate feeding behavior rapidly and without training. Nat Neurosci 14, 351–355 (2011).

3. M. J. Krashes et al., Rapid, reversible activation of AgRP neurons drives feeding behavior in mice. J Clin Invest 121, 1424–1428 (2011).

4. G. J. Morton, D. E. Cummings, D. G. Baskin, G. S. Barsh, M. W. Schwartz, Central nervous system control of food intake and body weight. Nature 443, 289–295 (2006).

5. M. L. Andermann, B. B. Lowell, Toward a Wiring Diagram Understanding of Appetite Control. Neuron 95, 757–778 (2017).

6. E. Roh, D. K. Song, M. S. Kim, Emerging role of the brain in the homeostatic regulation of energy and glucose metabolism. Exp Mol Med 48, e216 (2016).

7. L. Vong et al., Leptin action on GABAergic neurons prevents obesity and reduces inhibitory tone to POMC neurons. Neuron 71, 142–154 (2011).

8. M. H. Hastings, E. S. Maywood, M. Brancaccio, Generation of circadian rhythms in the suprachiasmatic nucleus. Nat Rev Neurosci 19, 453–469 (2018).

9. X. Peng, Y. Chen, The emerging role of circadian rhythms in the development and function of thermogenic fat. Front Endocrinol (Lausanne) 14, 1175845 (2023).

10. J. H. Strubbe, S. C. Woods, The timing of meals. Psychol Rev 111, 128–141 (2004).

11. H. P. Ostendorff et al., Ubiquitination-dependent cofactor exchange on LIM homeodomain transcription factors. Nature 416, 99–103 (2002).

12. H. P. Ostendorff et al., Dynamic expression of LIM cofactors in the developing mouse neural tube. Dev Dyn 235, 786–791 (2006).

13. B. Jiao et al., Functional activity of RLIM/Rnf12 is regulated by phosphorylation-dependent nucleocytoplasmic shuttling. Mol Biol Cell 24, 3085–3096 (2013).

14. F. Bustos et al., Functional Diversification of SRSF Protein Kinase to Control Ubiquitin-Dependent Neurodevelopmental Signaling. Dev Cell 55, 629–647 e627 (2020).

15. C. Gungor et al., Proteasomal selection of multiprotein complexes recruited by LIM homeodomain transcription factors. Proc Natl Acad Sci U S A 104, 15000–15005 (2007).

16. S. A. Johnsen et al., Regulation of estrogen-dependent transcription by the LIM cofactors CLIM and RLIM in breast cancer. Cancer Res 69, 128–136 (2009).

17. C. Gontan et al., RNF12 initiates X-chromosome inactivation by targeting REX1 for degradation. Nature 485, 386–390 (2012).

18. J. Shin et al., Maternal Rnf12/RLIM is required for imprinted X-chromosome inactivation in mice. Nature 467, 977–981 (2010).

19. F. Wang et al., Roles of the Rlim-Rex1 axis during X chromosome inactivation in mice. Proc Natl Acad Sci U S A 120, e2313200120 (2023).

20. F. Wang et al., Regulation of X-linked gene expression during early mouse development by Rlim. Elife 5, (2016).

21. J. Shin et al., RLIM is dispensable for X-chromosome inactivation in the mouse embryonic epiblast. Nature 511, 86–89 (2014).

22. F. Wang et al., Deficient spermiogenesis in mice lacking Rlim. Elife 10, (2021).

23. C. Contreras, R. Nogueiras, C. Dieguez, K. Rahmouni, M. Lopez, Traveling from the hypothalamus to the adipose tissue: The thermogenic pathway. Redox Biol 12, 854–863 (2017).

24. J. Wu et al., Beige adipocytes are a distinct type of thermogenic fat cell in mouse and human. Cell 150, 366–376 (2012).

25. B. Jiao et al., Paternal RLIM/Rnf12 is a survival factor for milk-producing alveolar cells. Cell 149, 630–641 (2012).

26. I. K. Franklin, C. B. Wollheim, GABA in the endocrine pancreas: its putative role as an islet cell paracrine-signalling molecule. J Gen Physiol 123, 185–190 (2004).

27. N. P. Hyland, J. F. Cryan, A Gut Feeling about GABA: Focus on GABA(B) Receptors. Front Pharmacol 1, 124 (2010).

28. M. M. Li et al., The Paraventricular Hypothalamus Regulates Satiety and Prevents Obesity via Two Genetically Distinct Circuits. Neuron 102, 653–667 e656 (2019).

29. J. Y. Lee et al., RIP-Cre revisited, evidence for impairments of pancreatic beta-cell function. J Biol Chem 281, 2649–2653 (2006).

30. Y. Hao et al., Integrated analysis of multimodal single-cell data. Cell 184, 3573–3587 e3529 (2021).

31. Y. Hashikawa et al., Transcriptional and Spatial Resolution of Cell Types in the Mammalian Habenula. Neuron 106, 743–758 e745 (2020).

32. L. Chung, A Brief Introduction to the Transduction of Neural Activity into Fos Signal. Dev Reprod 19, 61–67 (2015).

33. A. McDavid et al., Data exploration, quality control and testing in single-cell qPCR-based gene expression experiments. Bioinformatics 29, 461–467 (2013).

34. M. Y. Cheng et al., Prokineticin 2 transmits the behavioural circadian rhythm of the suprachiasmatic nucleus. Nature 417, 405–410 (2002).

35. H. M. Prosser et al., Prokineticin receptor 2 (Prokr2) is essential for the regulation of circadian behavior by the suprachiasmatic nuclei. Proc Natl Acad Sci U S A 104, 648–653 (2007).

36. S. J. Aton, C. S. Colwell, A. J. Harmar, J. Waschek, E. D. Herzog, Vasoactive intestinal polypeptide mediates circadian rhythmicity and synchrony in mammalian clock neurons. Nat Neurosci 8, 476–483 (2005).

37. E. S. Maywood, J. E. Chesham, J. A. O’Brien, M. H. Hastings, A diversity of paracrine signals sustains molecular circadian cycling in suprachiasmatic nucleus circuits. Proc Natl Acad Sci U S A 108, 14306–14311 (2011).

38. N. P. Achilly, Properties of VIP+ synapses in the suprachiasmatic nucleus highlight their role in circadian rhythm. J Neurophysiol 115, 2701–2704 (2016).

39. W. D. Todd et al., Suprachiasmatic VIP neurons are required for normal circadian rhythmicity and comprised of molecularly distinct subpopulations. Nat Commun 11, 4410 (2020).

40. S. Paul et al., Output from VIP cells of the mammalian central clock regulates daily physiological rhythms. Nat Commun 11, 1453 (2020).

41. J. C. Bruning, H. Fenselau, Integrative neurocircuits that control metabolism and food intake. Science 381, eabl7398 (2023).

42. D. A. M. Joye et al., Reduced VIP Expression Affects Circadian Clock Function in VIP-IRES-CRE Mice (JAX 010908). J Biol Rhythms 35, 340–352 (2020).

43. J. R. M. Harvey, A. E. Plante, A. L. Meredith, Ion Channels Controlling Circadian Rhythms in Suprachiasmatic Nucleus Excitability. Physiol Rev 100, 1415–1454 (2020).

44. E. E. Benarroch, HCN channels: function and clinical implications. Neurology 80, 304–310 (2013).

45. K. L. Eckel-Mahan et al., Reprogramming of the circadian clock by nutritional challenge. Cell 155, 1464–1478 (2013).

46. D. Kraus, Q. Yang, B. B. Kahn, Lipid Extraction from Mouse Feces. Bio Protoc 5, (2015).

47. C. Anaclet et al., The GABAergic parafacial zone is a medullary slow wave sleep-promoting center. Nat Neurosci 17, 1217–1224 (2014).

48. P. Gimenez-Gomez et al., Suppression of binge alcohol drinking by an inhibitory neuronal ensemble in the mouse medial orbitofrontal cortex. Nat Neurosci, (2025).

49. P. M. Klenowski et al., A neuronal coping mechanism linking stress-induced anxiety to motivation for reward. Sci Adv 9, eadh9620 (2023).

